# Deltex and RING-UIM E3 ligases cooperate to create a ubiquitin-ADP-ribose hybrid mark on tankyrase, promoting its stabilization

**DOI:** 10.1101/2025.04.09.648013

**Authors:** Jerome Perrard, Kevin Gao, Katherine Ring, Susan Smith

**Author notes:** These authors contributed equally (Jerome Perrard, Kevin Gao, Katherine Ring). Corresponding author Susan Smith.

## Abstract

ADP-ribosylation is a key post-translational modification that impacts diverse cellular pathways. The modification can occur as mono-ADP-ribose (MAR) or be extended into poly-ADP-ribose (PAR). Tankyrase, a PAR transferase, adds PAR to itself and other proteins to influence their function and stability by tagging them for proteasomal degradation via the PAR-binding E3 ligase RNF146. This degradation can be counteracted by RING-UIM E3 ligases RNF114 and RNF166, though the process is unclear. Here we identify a new mechanism that can regulate the balance between MAR and PAR on tankyrase to control degradation. We show that Deltex E3 ligases DTX2 and DTX3 catalyze monoubiquitylation of tankyrase in cells. This ubiquitylation occurs, not on a (canonical) lysine, but rather on mono-ADP-ribose, creating a monoubiquitin- MAR hybrid mark. RNF114 and RNF166 recognize this mark using a unique hybrid reader domain comprising two binding sites - one for MAR and one for ubiquitin - and further diubiquitylate it. This ubiquitylation of MAR, which occurs near the ADP-ribose addition site, prevents PAR formation, antagonizing the action of the PAR-binding E3 ligase RNF146 and stabilizing tankyrase. These findings reveal a novel interplay between ubiquitin, ADP-ribose, and E3 ligases in cellular signaling.

## Introduction

Post-translational modifications (PTMs) regulate protein function and affect various cellular processes. ADP-ribosylation is a key PTM that impacts all forms of life. It is catalyzed by the ADP-ribosyl-transferase (ART) enzyme superfamily^1, 2^, which transfers ADP-ribose from NAD^+^ onto amino acids in proteins including serine, threonine, tyrosine, glutamate, aspartate, and others^2^. This modification can remain as a single mono-ADP-ribose (MAR) or be extended into a poly-ADP-ribose (PAR) chain. MAR and PAR are recognized by reader domains, such as the macrodomain (which binds both MAR and the terminal ADP-ribose of PAR), and the WWE (tryptophan-tryptophan-glutamate) domain, which specifically reads PAR through iso-ADP-ribose, a molecule derived from two adjacent ADP-ribose units of PAR chain^2^.

Only four enzymes in the cell synthesize PAR chains: PARP1, PARP2, tankyrase 1, and tankyrase 2^3, 4^. These poly-ADP-ribosyltransferases (ARTs) belong to the seventeen-member PARP family, identified by their catalytic ART (also known as PARP) domains. While the remaining thirteen family members were expected to function as polyARTs, eleven are monoARTs and two are catalytically inactive^1, 5^. The activity of the four polyARTs can be reversed by the PAR-glycohydrolase (PARG), the main PAR-degrading enzyme in cells^6^, which efficiently targets PAR but is less effective on the last amino acid-bound MAR. The final MAR is cleaved by other enzymes like ARH3 and TARG, which cleave the last MAR, each targeting specific linkages such as serine and glutamate, respectively^6^.

Tankyrase 1 and 2 are related multifunctional proteins involved in various cellular pathways and linked to diseases, including cancer^3, 4, 7, 8^. Apart from their catalytic PARP domains, tankyrases are structurally distinct from PARP1 and 2. In addition to their C-terminal PARP domain, tankyrases contain a SAM domain that facilitates dimerization and an ankyrin repeat domain that recognizes binding motifs in diverse partners^3, 4^. Tankyrases PARylate themselves and specific binding partners. A key partner is the PAR-binding E3 ligase RNF146, which is activated by PAR to ubiquitylate and degrade tankyrase and its partners^9^.

Ubiquitylation, like ADP-ribosylation, is a significant PTM that regulates protein stability and function. Ubiquitin, a 76-amino acid protein, has a C-terminal glycine and seven lysine (K) residues^10^. Typically, ubiquitylation starts by linking ubiquitin’s C-terminal glycine to a lysine in the substrate. Additional ubiquitin molecules can be added through one of the seven lysines, forming a polyubiquitin chain. K48-linked chains are common for proteasomal degradation. This process requires enzyme E1 (activating), E2 (conjugating), and E3 (ligating) enzymes^11^, with E3s providing substrate specificity. RNF146 is a RING E3 ligase comprising a catalytic RING domain, a PAR-binding WWE domain, and tankyrase-binding motifs^12–16^. PAR-binding activates the RING domain to catalyze K48-linked polyubiquitylation of tankyrases and subsequent proteasomal degradation^9^.

We recently discovered that RNF146-mediated degradation of tankyrase can be countered by RING-UIM E3 ligases RNF114 and RNF166^17^. RING-UIMs have an N-terminal RING domain, a C2H2 zinc finger (ZnF), a Di19 domain with two ZnFs (so named because it resembles plant Di19 gene family domains), and a C-terminal ubiquitin interacting motif (UIM)^18, 19^. We identified a monoubiquitylated form of tankyrase (monoUbTNKS) in cells that is a target for RNF114/166. We found that the C-terminal Di19-UIM fragment of RNF166 binds monoUbTNKS, and the N-terminal RING catalyzes K11-linked diubiquitylation, which competes with RNF146- mediated K48-linked polyubiquitylation and degradation and stabilizes tankyrase^17^. Tankyrase binding to RNF114/166 depends on its catalytic activity^17^, but RNF114/166 lack WWE domains or other known PAR-binding motifs. A recent study identified a MAR binding domain in RNF114^20^. Using specific recombinant antibodies against MAR, we showed that cellular tankyrase is modified by MAR in addition to PAR^17^, providing a potential mechanism for RNF114/166 binding to tankyrase. The E3 ligase that generates the RNF114/166 target (monoUbTNKS) is not known.

Recent studies have linked the Deltex family of E3 ligases (DTX1, 2, 3, 3L, and 4)^21, 22^ to ADP-ribosylation. All Deltex E3s have a catalytic RING and C-terminal domain (CTD) containing an ADP-ribose-binding site that recruits PARP1/2-mediated ADP-ribosylated proteins for ubiquitylation^23^. DTX1, 2, and 4 exclusively have PAR-binding WWE domains at their N-termini and bind tankyrase, dependent on its catalytic activity^17^, but the physiological relevance is not known. Recent studies showed that Deltex E3 ligases can catalyze ubiquitylation of ADP-ribose on peptides and proteins in vitro creating a novel protein-ADP-ribose-ubiquitin linkage^24^.

Whether DTX E3s catalyze this reaction in cells is not known.

We aimed to characterize the origin and impact of MARylated, PARylated, monoUbTNKS in cells. We found that DTX2 and DTX3 catalyze monoubiquitylation of MAR on tankyrase, forming a unique hybrid mark that is bound and stabilized by a hybrid reader in RNF114/166 and further diubiquitylated. The ubiquitylation of MAR prevents PAR formation and consequent degradation by RNF146, thereby stabilizing tankyrase. Our findings reveal crosstalk between ubiquitin and ADP-ribose, with broad implications for cellular regulation.

## Results

### Tankyrase is ubiquitylated on MAR

We previously identified a monoubiquitylated form of tankyrase (monoUbTNKS) in cells, which is a target for ubiquitylation by the RNF114 or RNF166 RING-UIM E3 ligases^17^. In our previous work, we described how the C-terminal Di19-UIM fragment of RNF166 binds and stabilizes monoUbTNKS, while the N-terminal RING catalyzes K11-linked diubiquitylation (Fig. 1A). To further investigate the site of ubiquitin attachment, we isolated monoUbTNKS from cells using the SAMPARP fragment of tankyrase 1. We chose SAMPARP rather than full-length tankyrase because it was sufficient for both the mono- and diubiquitylation and its smaller size (55 kDa versus 135 kDa) made it easier to resolve on SDS-PAGE. We introduced Myc-tagged SAMPARP, HA-tagged ubiquitin (HA-Ub), and Flag-tagged Di19-UIM (from RNF166) into HEK293T cells deleted for tankyrases (TNKS1/2 DKO cells^25^). The HA-ubiquitylated proteins were isolated using anti-HA antibody affinity matrix (HA-Beads) and detected with anti-Myc antibody. As shown in Fig. 1B (lane 6) we observed monoubiquitylated MycSAMPARP, which required all three components. We also detected unmodified MycSAMPARP, likely due to its multimerization with HA-UbMycSAMPARP or degradation of the ubiquitin modification during isolation. Probing the anti-HA immunoprecipitated proteins with anti-HA antibody confirmed the identity of the HA-monoubiquitylated species (extra panel, lane 6).

**Figure 1.**
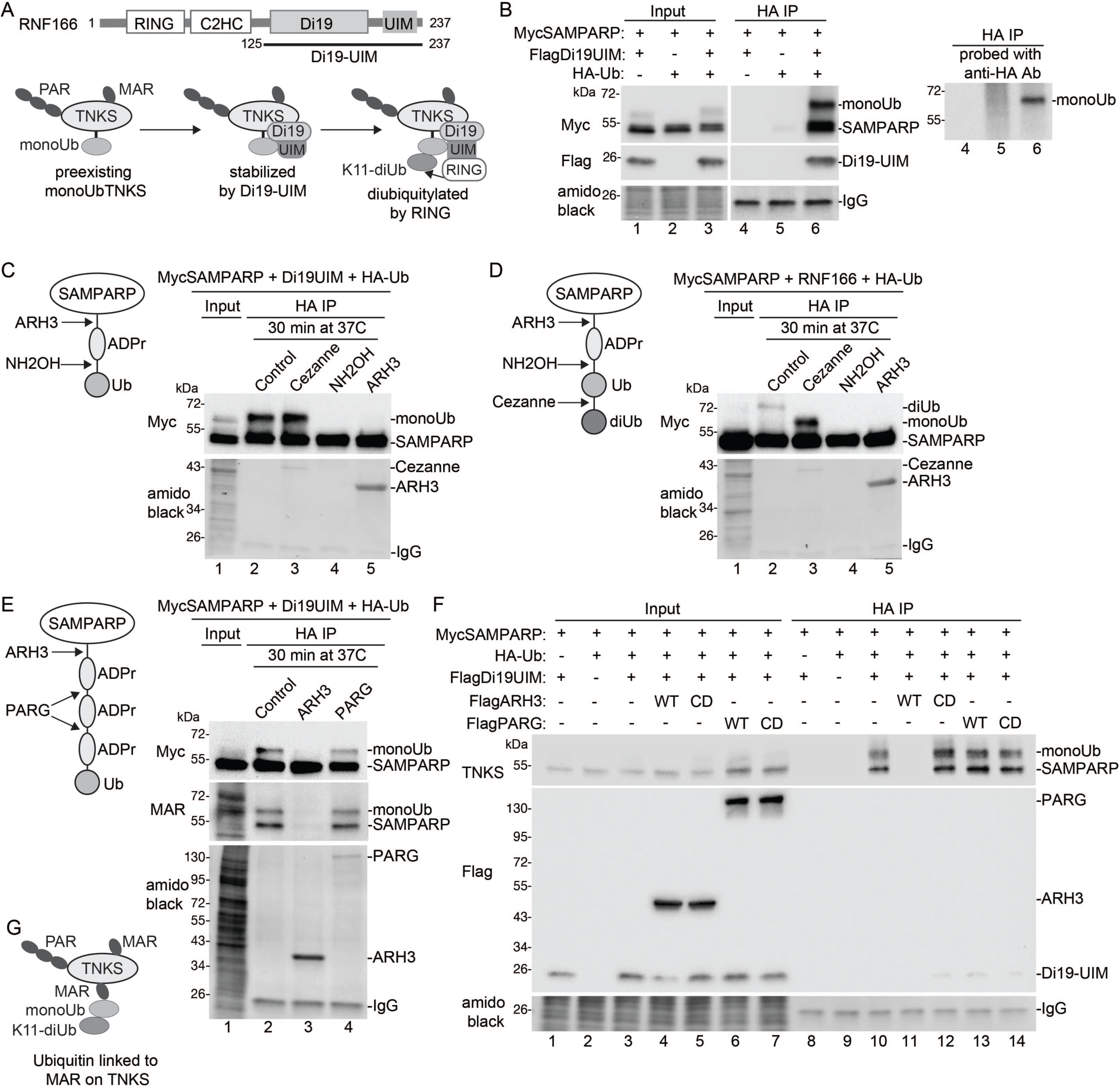
TNKS is ubiquitylated on mono-ADP-ribose. A. Schematic diagram of RNF166 and the Di19UIM construct. The pathway for RNF166- mediated K11-linked diubiquitylation of TNKS is shown below. B. Immunoblot analysis of TNKS1/2 DKO HEK293T cells transfected with MycSAMPARP, FlagDi19UIM and HA-Ub plasmids, immunoprecipitated with anti-HA antibody, stained with amido black, and probed with antibodies indicated on the left. Right panel shows HA IP reprobed with anti-HA antibody. C. Schematic diagram showing the cleavage sites for ARH3 and NH2OH on Ub-MAR-SAMPARP. Immunoblot analysis of Ub-MAR-MycSAMPARP that was HA-immuno-isolated from TNKS1/2 DKO HEK293T cells transfected with MycSAMPARP, FlagD19UIM, and HA-Ub plasmids, and then incubated in vitro for 30 min at 37°C with the indicated treatments, stained with amido black, and probed with anti-Myc antibody. D. Schematic diagram showing the cleavage sites for ARH3, NH2OH, and Cezanne on diUb-MAR- SAMPARP. Immunoblot analysis of diUb-MAR-MycSAMPARP that was HA-immuno-isolated from TNKS1/2 DKO HEK293T cells transfected with MycSAMPARP, FlagRNF166, and HA-Ub plasmids, and then incubated in vitro for 30 min at 37°C with the indicated treatments, stained with amido black, and probed with anti-Myc antibody. E. Schematic diagram showing the cleavage sites for ARH3 and PARG on Ub-PAR-MycSAMPARP. Immunoblot analysis of Ub-MAR-MycSAMPARP that was HA-immuno-isolated from denatured TNKS1/2 DKO HEK293T cells transfected with the MycSAMPARP, FlagD19UIM, and HA-Ub plasmids, and then incubated in vitro for 30 min at 37°C with the indicated treatments, stained with amido black, and probed with antibodies indicated on the left. F. Immunoblot analysis of TNKS1/2 DKO HEK293T cells transfected with MycSAMPARP, HA-Ub, and the indicated Flag plasmids, immunoprecipitated with anti-HA antibody, stained with amido black, and probed with the antibodies indicated on the left. G. Schematic diagram showing K11-linked diubiquitin on MAR on TNKS.

We initially assumed that monoubiquitin on TNKS was attached to a canonical lysine.

After tryptic digestion and mass spectrometry, we detected TNKS but not ubiquitylated peptides. Considering recent studies showing ADP-ribose-linked ubiquitylation^24^, we hypothesized that monoubiquitin on SAMPARP might be linked via ADP-ribose, rather than lysine (see schematic prediction Fig. 1C). MonoUbSAMPARP was isolated from transfected cells as in Fig. 1B and incubated in vitro. The control (Fig. 1C, lane 2) showed monoUbSAMPARP. Treatment with Cezanne, a K11-linkage-specific deubiquitinase^26, 27^, which we showed previously cleaved the diubiquitin but not the monoubiquitin off SAMPARP^17^ had no effect (lane 3), as expected. Treatment with hydroxylamine (NH2OH), which cleaves ester and phosphoanhydride but not peptidic or glycosidic bonds (and thus could cleave bonds between ADP-ribose and ubiquitin or within ADP-ribose), eliminated monoUb on SAMPARP (lane 4), consistent with a ubiquitin-ADP-ribose linkage. Treatment with ARH3, a glycohydrolase that cleaves the bond between ADP-ribose and the amino acid (usually serine), also cleaved the monoubiquitin off SAMPARP (lane 5). These results indicate that monoubiquitin is likely linked to a serine on SAMPARP via ADP-ribose, as shown in the schematic (Fig. 1C).

We then investigated whether the K11-linked diubiquitin on SAMPARP was linked to the monoubiquitin linked to ADP-ribose (see schematic prediction, Fig. 1D). DiUbSAMPARP was isolated from transfected cells as in Fig. 1B (except full length FlagRNF166 was used instead of FlagDi19-UIM) and incubated in vitro. The control (Fig. 1D, lane 2) showed diUbSAMPARP. Cezanne cleaved the diubiquitin, leaving monoUbSAMPARP (lane 3), as expected. Treatment with NH2OH (lane 4) or ARH3 (lane 5) removed the diubiquitin, indicating that the K11-linked diubiquitin is attached to the monoubiquitin on ADP-ribose on SAMPARP, as shown in the schematic (Fig. 1D).

We investigated whether monoubiquitin is linked to MAR or PAR on SAMPARP (see schematic prediction, Fig. 1E). MonoUbSAMPARP was isolated from transfected cells as in Fig. 1B and incubated in vitro. The control (Fig. 1E, lane 2) showed monoUbSAMPARP. Treatment with ARH3 cleaved the monoubiquitin off SAMPARP (lane 3), while PARG had no effect (lane 4). PARG degrades PAR by cleaving between ADP-ribose units, but unlike ARH3, is inefficient at cleaving the bond between ADP-ribose and the amino acid^6^. Immunoblotting with a MAR-specific antibody (Fig. 1E, middle panel) revealed that both SAMPARP and monoUbSAMPARP were MARylated (lane 2). ARH3 removed both MAR and ubiquitin (lane 3), while PARG did not (lane 4). These results together indicate that monoubiquitin is not on PAR as shown schematically in Fig. 1E, rather monoubiquitin is on MAR (shown schematically in Fig. 1C and D).

To confirm the in vitro results, we performed the same analysis in cells. MycSAMPARP, HA-Ub, and FlagDi19-UIM were transfected into TNKS1/2 DKO cells with FlagARH3 (WT or CD; a catalytically dead mutant D77/78N^28^) or FlagPARG (WT or CD; a catalytically dead mutant E755/756A^29^). HA-ubiquitylated proteins were immunoisolated and analyzed by immunoblot. As shown in Fig. 1F, monoUbSAMPARP (lane 10) was removed by ARH3 WT (lane 11), but not CD (lane 12), and was unaffected by PARG WT (lane 13) or CD (lane 14). These results confirm that tankyrase is ubiquitylated on MAR (shown schematically in Fig. 1G).

### DTX2 monoubiquitylates MAR on tankyrase

We aimed to identify the E3 ligase responsible for monoubiquitylation of MARylated TNKS. Since induction required TNKS catalytic activity^17^, we initially hypothesized that a PAR-binding E3 ligase like RNF146 might be involved. However, this was ruled out, as Di19UIM induced monoUbTNKS in RNF146 KO cells^17^. We then considered other E3 ligases with PAR-binding WWE domains, including three members of the Deltex family (DTX1, 2, and 4) (Fig. 2A), which bind TNKS dependent on its catalytic activity^17^. DTX2 was previously shown to ubiquitylate PARP1/2-mediated ADP-ribosylated proteins in vitro^24^. We focused on DTX2 as mass spectrometry data indicated that it was the only member of the family expressed at detectable levels in HEK293T cells^30^, making it a likely candidate responsible for generating monoUbMARTNKS.

**Figure 2.**
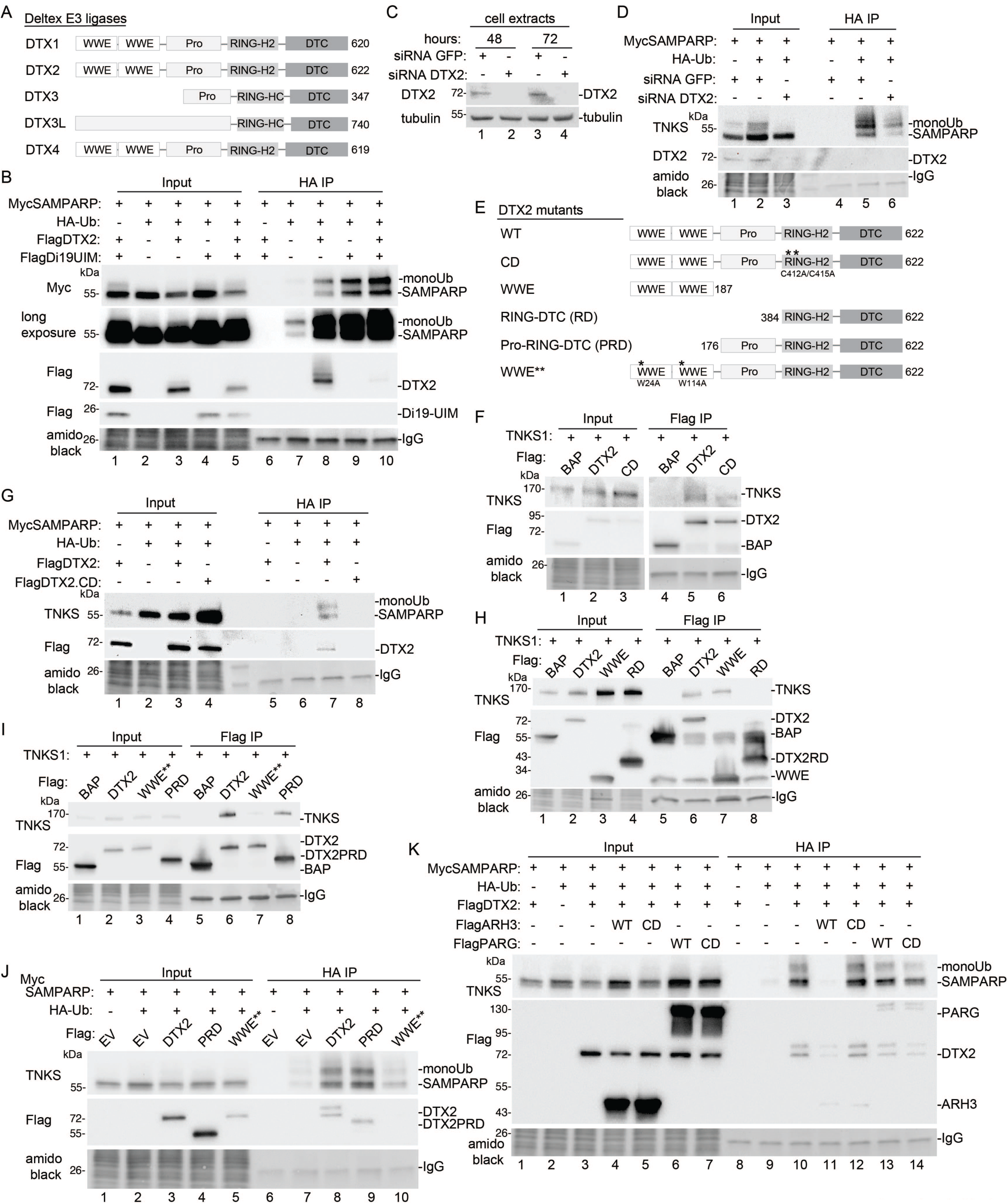
DTX2 monoubiquitylates TNKS on mono-ADP-ribose. A. Schematic diagram showing the five members of the Deltex E3 ligase family. B. Immunoblot analysis of TNKS1/2 DKO HEK293T cells transfected with MycSAMPARP, HA-Ub and the indicated Flag plasmids, immunoprecipitated from denatured cell extracts with anti-HA antibody, stained with amido black, and probed with antibodies indicated on the left. C. Immunoblot analysis of TNKS1/2 DKO HEK293T cells transfected with GFP or DTX2 siRNA and probed with antibodies indicated on the left. D. Immunoblot analysis of TNKS1/2 DKO HEK293T cells transfected with MycSAMPARP, HA-Ub and GFP or DTX2 siRNA, immunoprecipitated with anti-HA antibody, stained with amido black, and probed with anti-Myc-antibody. E. Schematic diagram showing DTX2 mutant constructs. F. Immunoblot analysis of TNKS1/2 DKO HEK293T cells transfected with TNKS1and the indicated Flag plasmids, immunoprecipitated with anti-Flag antibody, stained with amido black, and probed with antibodies indicated on the left. G. Immunoblot analysis of TNKS1/2 DKO HEK293T cells transfected with the MycSAMPARP, HA-Ub and the indicated Flag plasmids, immunoprecipitated with anti-HA antibody, stained with amido black, and probed with antibodies indicated on the left. H. Immunoblot analysis of TNKS1/2 DKO HEK293T cells transfected with TNKS1and the indicated Flag plasmids, immunoprecipitated with anti-Flag antibody, stained with amido black, and probed with antibodies indicated on the left. I. Immunoblot analysis of TNKS1/2 DKO HEK293T cells transfected with TNKS1and the indicated Flag plasmids, immunoprecipitated with anti-Flag antibody, stained with amido black, and probed with antibodies indicated on the left. J. Immunoblot analysis of TNKS1/2 DKO HEK293T cells transfected with the MycSAMPARP, HA- Ub and the indicated Flag plasmids, immunoprecipitated with anti-HA antibody, stained with amido black, and probed with antibodies indicated on the left. K. Immunoblot analysis of TNKS1/2 DKO HEK293T cells transfected with the MycSAMPARP, HA-Ub and the indicated Flag plasmids, immunoprecipitated with anti-HA antibody, stained with amido black, and probed with antibodies indicated on the left.

We overexpressed DTX2 to test if it catalyzed monoubiquitylation of TNKS. TNKS1/2 DKO cells were transfected with DTX2, MycSAMPARP, and HA-ubiquitin. The HA-ubiquitylated proteins were isolated using HA-Beads and detected by immunoblot with anti-Myc antibody. As shown in Fig. 2B, lane 8, DTX2 promoted monoubiquitylation of SAMPARP. A similar product was seen with the RNF166 Di19-UIM fragment alone (lane 9) and combining DTX2 with Di19-UIM further increased monoubiquitylation (lane 10). A longer exposure revealed monoUbSAMPARP even without DTX2 or Di19-UIM (lane 7), likely due to endogenous DTX2.

To test if endogenous DTX2 is responsible for monoubiquitylation of TNKS SAMPARP, we depleted it using siRNA. TNKS1/2 DKO cells were transfected with GFP (control) or DTX2 siRNA and analyzed by immunoblot after 48 or 72 hours. Fig. 2C shows endogenous DTX2 was present in control cells (lanes 1 and 3) and depleted with DTX2 siRNA (lanes 2 and 4). For ubiquitylation analysis, TNKS1/2 DKO cells were treated with GFP or DTX2 siRNA and transfected with MycSAMPARP and HA-ubiquitin. HA-ubiquitylated proteins were isolated using HA-Beads and detected by immunoblot with anti-TNKS antibody. As shown in Fig. 2D, DTX2 siRNA led to DTX2 depletion (Input, lane 3) and reduced monoubiquitylation of SAMPARP (IP, lane 6).

To further characterize the reaction, we created DTX2 mutants (Fig. 2E) and measured TNKS binding (via Flag IP) and monoubiquitylation (via isolation with HA-Beads). To test if E3 ligase activity was required, we generated a catalytically dead (CD, C412/415A) RING mutant^31^. FLAG IP showed that both WT and CD DTX2 bound TNKS (Fig. 2F, lanes 5 and 6), though binding was slightly weaker for the CD mutant. HA IP revealed that only WT DTX2 catalyzed monoubiquitylation of TNKS SAMPARP (Fig. 2G, lane 7 versus lane 8). Next, we identified the TNKS-binding domains in DTX2. Flag IP (Fig. 2H) showed that the N-terminal WWE domains were sufficient for binding (lane 7), but a RING-DTC (RD) construct failed to bind TNKS (lane 8).

Mutating the essential amino acids in the WWE domains (WWE**, W24/114A)^14, 23^ in full length DTX2 led to reduced binding (Fig. 2I, lane 7 versus lane 6), suggesting that the WWE domains might be required. However, deletion of the WWE domains to generate a Pro-RING-DTC (PRD) construct, bound TNKS (lane 8). HA IP (Fig. 2J) showed that DTX2 PRD (lane 9), but not WWE** (lane 10), induced monoubiquitylation like WT DTX2 (lane 8). Together results demonstrate that the PRD fragment of DTX2 is sufficient to bind and monoubiquitylate TNKS SAMPARP.

To determine if the DTX2-mediated monoUb is on MAR, we transfected TNKS1/2 DKO cells with DTX2, MycSAMPARP, HA-Ub, along with FlagARH3 (WT or CD) or FlagPARG (WT or CD). HA-ubiquitylated proteins were isolated using HA-Beads and detected by immunoblot with anti-TNKS antibody. As shown in Fig. 2K, monoUbSAMPARP (lane 10) was removed by ARH3 WT (lane 11), but not CD (lane 12) and not by PARG WT (lane 13) or CD (lane 14). These results indicate that DTX2 catalyzes monoubiquitylation of MAR, likely on a serine residue of TNKS SAMPARP.

### DTX3 monoubiquitylates MAR on tankyrase

Our results show that the WWE domains are not necessary for DTX2 binding and monoubiquitylation of TNKS, so we examined other Deltex family members, DTX3 and DTX3L, which lack WWE domains (Fig. 3A). As shown in Fig. 3B, DTX3 (lane 6), but not DTX3L (lane 7), co-immunoprecipitated TNKS and binding depended on TNKS catalytic activity (Fig. 3C, lanes 5 and 6). To assess whether DTX3 monoubiquitylates TNKS, we introduced DTX3, MycSAMPARP, and HA-ubiquitin into TNKS1/2 DKO cells. HA-ubiquitylated proteins were isolated using HA-Beads and detected by immunoblot with anti-TNKS antibody. As shown in Fig. 3D, lane 7, DTX3 monoubiquitylated SAMPARP. We also confirmed that DTX3’s catalytic activity was essential for ubiquitylation; a catalytically dead (CD, C164/167S^31^) RING mutant, which binds TNKS in an immunoprecipitation assay (Fig. 3E, lane 6), did not induce ubiquitylation of SAMPARP (Fig. 3D, lane 8).

**Figure 3.**
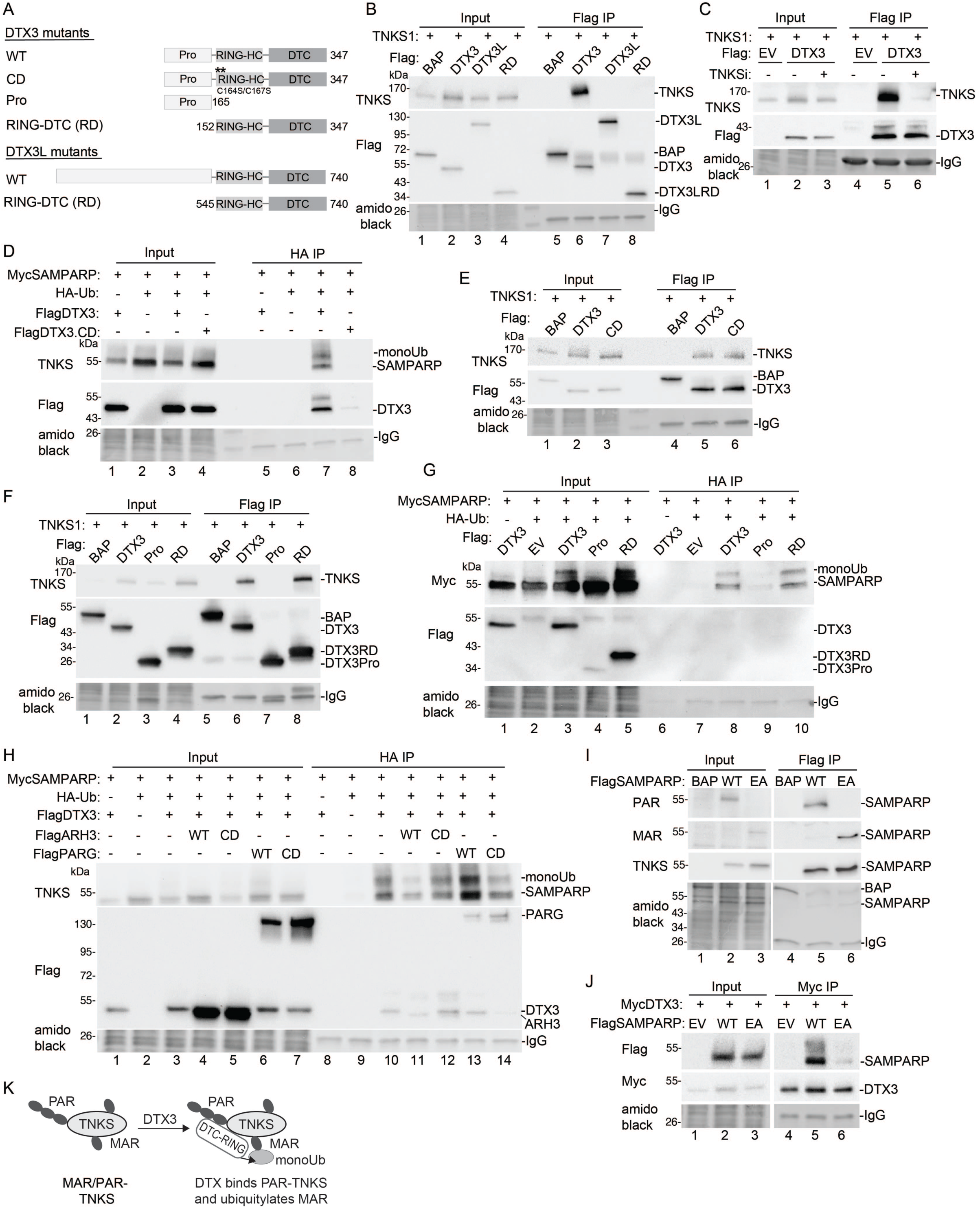
DTX3 monoubiquitylates TNKS on mono-ADP-ribose. A. Schematic diagram showing DTX3 and DTX3L mutant constructs. B. Immunoblot analysis of TNKS1/2 DKO HEK293T cells transfected with TNKS1and the indicated Flag plasmids, immunoprecipitated with anti-Flag antibody, stained with amido black, and probed with antibodies indicated on the left. C. Immunoblot analysis of TNKS1/2 DKO HEK293T cells transfected with TNKS1and the indicated Flag plasmids, with or without TNKSi, immunoprecipitated with anti-Flag antibody, stained with amido black, and probed with antibodies indicated on the left. D. Immunoblot analysis of TNKS1/2 DKO HEK293T cells transfected with the MycSAMPARP, HA-Ub and the indicated Flag plasmids, immunoprecipitated with anti-HA antibody, stained with amido black, and probed with antibodies indicated on the left. E. Immunoblot analysis of TNKS1/2 DKO HEK293T cells transfected with TNKS1and the indicated Flag plasmids, immunoprecipitated with anti-Flag antibody, stained with amido black, and probed with antibodies indicated on the left. F. Immunoblot analysis of TNKS1/2 DKO HEK293T cells transfected with TNKS1and the indicated Flag plasmids, immunoprecipitated with anti-Flag antibody, stained with amido black, and probed with antibodies indicated on the left. G. Immunoblot analysis of TNKS1/2 DKO HEK293T cells transfected with the MycSAMPARP, HA-Ub and the indicated Flag plasmids, lysed under denaturing conditions, immunoprecipitated with anti-HA antibody, stained with amido black, and probed with antibodies indicated on the left. H. Immunoblot analysis of TNKS1/2 DKO HEK293T cells transfected with the MycSAMPARP, HA- Ub and the indicated Flag plasmids, immunoprecipitated with anti-HA antibody, stained with amido black, and probed with antibodies indicated on the left. I. Immunoblot analysis of TNKS1/2 DKO HEK293T cells transfected with the FlagBAP or FlagSAMPARP (WT or EA), immunoprecipitated with anti-Flag antibody, stained with amido black, and probed with antibodies indicated on the left. J. Immunoblot analysis of TNKS1/2 DKO HEK293T cells transfected with the MycSAMPARP, and FlagSAMPARP (WT or EA), immunoprecipitated with anti-Myc antibody, stained with amido black, and probed with antibodies indicated on the left. K. Schematic diagram showing DTX3 (RD) binds PAR and ubiquitylates MAR on TNKS.

To map the domains of DTX3 involved in binding and ubiquitylation, we created Pro and RING-DTC (RD) constructs. The DTX3 RD domain was both necessary and sufficient for binding (Fig. 3F, lane 8) and monoubiquitylation (Fig. 3G, lane 10) of TNKS. Despite the high degree of homology between the DTX3 and DTX3L RD domains, DTX3L did not bind TNKS (see Fig. 3B, lane7). We hypothesized that the large N-terminal domain of DTX3L might be inhibitory, but deleting this domain to create a DTX3L RD construct (Fig. 3A) did not reveal an interaction with TNKS (Fig. 3B, lane 8).

To determine if the DTX3-mediated monoubiquitylation of SAMPARP is on MAR, we transfected DTX3, MycSAMPARP, and HA-Ub into TNKS1/2 DKO cells along with FlagARH3 (WT or CD) or FlagPARG (WT or CD), immunoisolated HA-ubiquitylated proteins, and detected HA- ubiquitylated MycSAMPARP by immunoblotting with anti-TNKS antibody. As shown in Fig. 3H, monoUbSAMPARP (lane 10) was reduced by ARH3 WT (lane 11), but not CD (lane 12) and not by PARG WT (lane 13) or CD (lane 14), indicating that DTX3 catalyzes monoubiquitylation of MAR likely on a serine residue of TNKS SAMPARP.

Our experiments show that WWE is not required for DTX-mediated binding and ubiquitylation of TNKS. While WWE binds PAR, the DTC domain can bind to MAR or PAR. To test if MARylated TNKS is sufficient for DTX binding, we introduced a point mutation (E1291A^32^) into the TNKS SAMPARP domain that eliminates PARylation but not MARylation. Immunoblotting confirmed this (Fig. 3I); FlagSAMPARP WT was detected by anti-PAR (and weakly by anti-MAR) antibody (lane 5), while the EA mutant was detected only by anti-MAR (lane 6) antibody.

MycDTX3 coimmunoprecipitated FlagSAMPARP WT (Fig. 3J, lane 5), but not the EA mutant (lane 6), indicating that MARylation alone is not sufficient for DTX3 binding. Overall, our data show that DTX3 binds PARylated TNKS and monoubiquitylates MAR (schematic representation Fig. 3K).

### Di19-UIM is a hybrid reader of the ubiquitin-MAR mark on tankyrase

What is the impact of the ubiquitin-MAR mark on TNKS? Ubiquitylation of MAR occurs at the 3′ hydroxyl of the adenine-proximal ribose ring of ADPribose^24^ adjacent to the 2’hydroxyl acceptor site for ADP-ribose polymer elongation (Fig. 4A). We hypothesized that ubiquitylation at this 3’ site, along with binding of Di19-UIM, could hinder ADP-ribose addition at the 2’ site, stabilizing MAR by preventing its extension to PAR. To test this, we induced monoubiquitylation of MAR on TNKS using Di19-UIM and measured MAR and PAR levels. TNKS1/2 DKO cells were transfected with FlagTNKS with or without Di19-UIM, immunoprecipitated with anti-Flag antibody, and analyzed by immunoblot. As shown in Fig. 4B, lane 5, both MARylated and PARylated TNKS were detected. Treatment with TNKSi (lane 4), abolished both signals, confirming dependence on TNKS activity. Cotransfection with Di19-UIM led to a marked increase in MARTNKS and a decrease in PARTNKS, relative to TNKS (lane 6).

**Figure 4.**
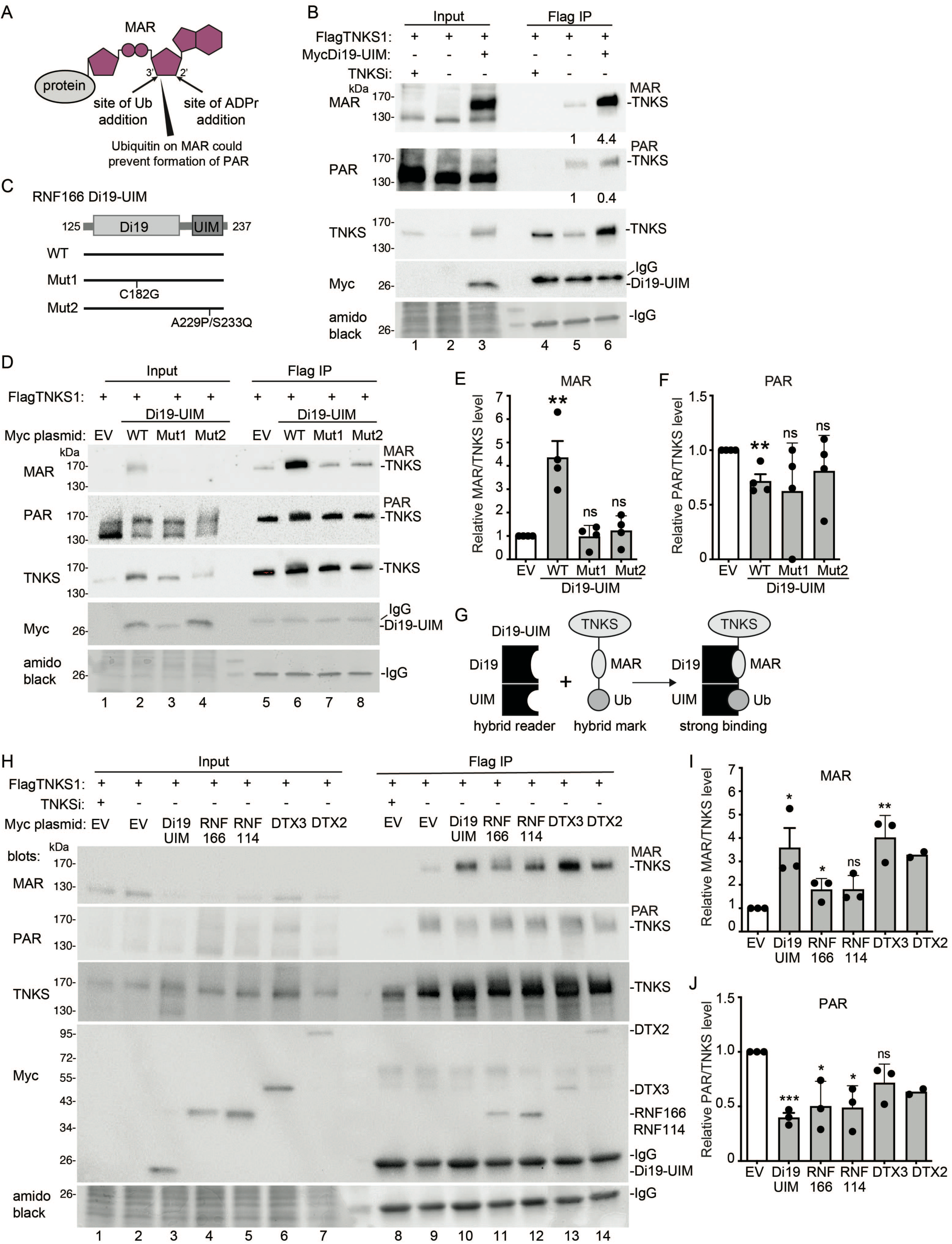
Di19-UIM is a hybrid reader of the ubiquitin-MAR mark on tankyrase. A. Schematic diagram showing MAR on protein. The sites of ubiquitin (Ub) addition and ADP-ribose (ADPr) addition on MAR are indicated. B. Immunoblot analysis of TNKS1/2 DKO HEK293T cells transfected with FlagTNKS1and MycDi19UIM, with or without TNKSi, immunoprecipitated with anti-Flag antibody, stained with amido black, and probed with antibodies indicated on the left. Protein levels relative to TNKS and normalized to the control are indicated below the blots. C. Schematic diagram of RNF166 and Di19-UIM WT and mutant constructs. D. Immunoblot analysis of TNKS1/2 DKO HEK293T cells transfected with FlagTNKS1and MycDi19UIM (WT, Mut1, or Mut2), immunoprecipitated with anti-Flag antibody, stained with amido black, and probed with antibodies indicated on the left. E. Graphical presentation of the change in MAR relative to TNKS1 induced by Di19UIM (WT, Mut1, or Mut2) and normalized EV. Average of four independent experiments ± SEM. Di19UIM WT vs EV: p = .003. Mut1 vs EV: p = 1.0. Mut2 vs EV: 0.5. **p ≤ 0.01, Student’s unpaired two- tailed t-test. ns, not significant. F. Graphical presentation of the change in PAR relative to TNKS1 induced by Di19UIM (WT, Mut1, or Mut2) and normalized to EV. Average of four independent experiments ± SEM. Di19UIM WT vs EV: p = .004. Mut1 vs EV: p = .14. Mut vs EV: p = 0.3. **p ≤ 0.01, Student’s unpaired two-tailed t-test. ns, not significant. G. Schematic diagram showing the Di19-UIM hybrid reader binding to the MAR-Ub hybrid mark on TNKS. H. Immunoblot analysis of TNKS1/2 DKO HEK293T cells transfected with FlagTNKS1and the indicated Myc plasmids, with or without TNKSi, immunoprecipitated with anti-Flag antibody, stained with amido black, and probed with antibodies indicated on the left. I. Graphical presentation of the change in MAR relative to TNKS1 induced by Di19UIM, RNF166, RNF144, DTX3, or DTX2, and normalized to EV. Average of three independent experiments ± SEM. Di19UIM vs EV: p = .04, RNF166 vs EV: p=.04, RNF114 vs EV: p=0.7 DTX3 vs EV: p= .005. *p ≤ 0.05, **p ≤ 0.01, Student’s unpaired two-tailed t-test. ns, not significant. J. Graphical presentation of the change in PAR relative to TNKS1 induced by Di19UIM, RNF166, RNF144, DTX3, or DTX2, and normalized to EV. Average of three independent experiments ± SEM. Di19UIM vs EV: p = .0001, RNF166 vs EV: p=.02, RNF114 vs EV: p=0.01, DTX3 vs EV: p= .07. *p ≤ 0.05, ***p ≤ 0.001, Student’s unpaired two-tailed t-test. ns, not significant.

The Di19-UIM fragment of RNF166 has two domains involved in TNKS interaction: the ZnFs in the Di19 domain and the ubiquitin-binding site in the UIM^17^. We aimed to determine the impact of each domain on MARTNKS induction. For Di19, we previously showed that mutations in the conserved cysteines of the first (C152R) or second (C182G) ZnF eliminated RNF166 binding to TNKS^17^. A recent study showed that deleting either ZnF or mutating the second ZnF’s conserved cysteine (C176A) in RNF114 Di19 prevented binding to a MARylated peptide^20^.

Based on this, we introduced the C182G mutation in the RNF166 Di19UIM construct (Fig. 4C; Mut1). For the UIM, we previously showed that a double mutation (A229P/S233Q) in the ubiquitin binding site reduced RNF166-mediated TNKS stabilization^17^. We introduced this mutation into the Di19UIM construct (Fig. 4C; Mut2).

We transfected TNKS1/2 DKO cells with FlagTNKS alongside MycDi19-UIM WT, Mut1, or Mut2, immunoprecipitated FlagTNKS, and performed immunoblot analysis. As shown in Fig. 4D, (and quantified in Fig. 4E and F), Di19-UIM WT caused a significant increase in MAR (lane 6).

This effect was abolished by mutation in the ZnF (Mut 1, lane 7) or the UIM (Mut2, lane 8), indicating both domains are required for MARTNKS induction. We propose that Di19-UIM acts as a hybrid reader for the UbMAR mark on TNKS (Fig. 4G). While the separate domains may provide weak interactions, combined they bind with high affinity and specificity. We hypothesize that binding stabilizes ubiquitylated MARylated TNKS by preventing extension of MAR to PAR. Finally, we tested the ability of full-length RNF166 and RNF114 (which diubiquitylate UbMARTNKS) and DTX2 and DTX3 (which monoubiquitylate MARTNKS) to induce MAR. As shown in Fig. 4H and quantified in Fig. I and J, each induced MARTNKS and reduced PARTNKS.

### Deltex and RING-UIM E3 ligases counter RNF146-mediated degradation of TNKS

RNF146-mediated degradation is crucial for regulating tankyrase stability and its activation depends on the PAR modification of TNKS^9^. We hypothesized that the Di19-UIM-induced reduction in PARylation could limit PAR-dependent RNF146-mediated degradation of TNKS. We transfected FlagTNKS and MycRNF146 into TNKS1/2 DKO cells and examined the effects of Di19-UIM and DTX3. As shown in Fig. 5A, top panel, RNF146 induced degradation of TNKS (lane 2), as expected. Di19-UIM had minimal effect (lane 3), while DTX3 increased TNKS (lane 4), slightly above control (lane 1). Combined Di19-UIM and DTX3 caused a dramatic increase in TNKS (lane 5). Anti-MAR detection (second panel) showed increased MARTNKS in lanes 4 and 5. Interesting, RNF146 protein levels (third panel) also increased in lanes 4 and 5. Since PAR stimulates RNF146 autoubiquitylation and degradation^12^, the increased MAR and reduced PAR from the DTX3/RNF166 pathway may limit autoubiquitylation and degradation of RNF146.

**Figure 5.**
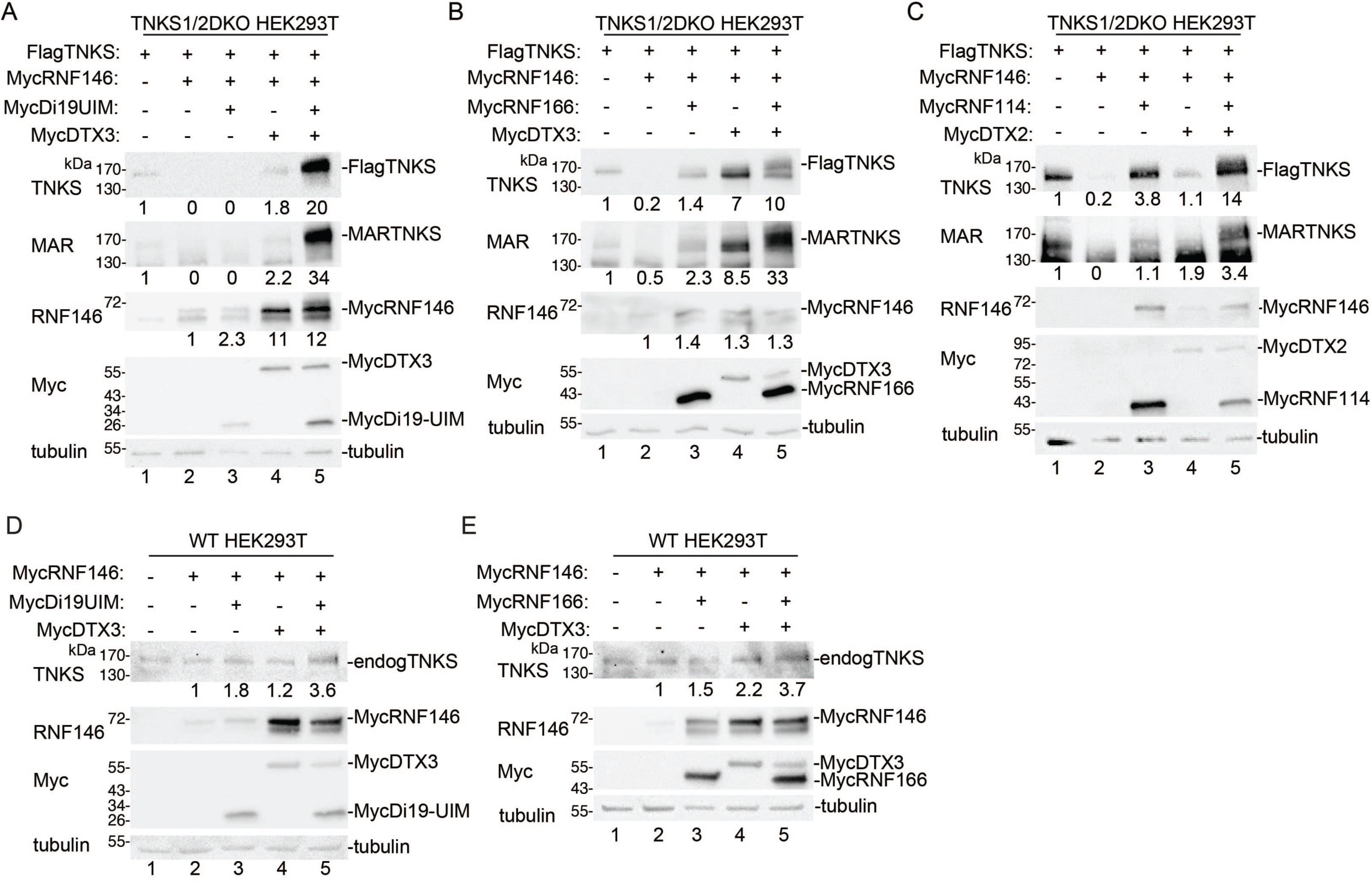
Deltex and RINGUIM E3 ligases counter RNF146-mediated degradation of TNKS. A. Immunoblot analysis of extracts from TNKS1/2 DKO HEK293T cells transfected with FlagTNKS1 and the indicated Myc plasmids, probed with antibodies indicated on the left. B. Immunoblot analysis of extracts from TNKS1/2 DKO HEK293T cells transfected with FlagTNKS1 and the indicated Myc plasmids, probed with antibodies indicated on the left. C. Immunoblot analysis of extracts from TNKS1/2 DKO HEK293T cells transfected with FlagTNKS1 and the indicated Myc plasmids, probed with antibodies indicated on the left. D. Immunoblot analysis of extracts from WT HEK293T cells transfected with the indicated Myc plasmids, probed with antibodies indicated on the left. E. Immunoblot analysis of extracts from WT HEK293T cells transfected with the indicated Myc plasmids, probed with antibodies indicated on the left.

We performed a similar analysis using full length RNF166 instead of Di19-UIM. As shown in Fig. 5B (top panel) RNF146 induced degradation of TNKS (lane 2). Addition of RNF166 or DTX3 increased TNKS levels (lanes 3 and 4), with combination of both leading to a greater increase (lane 5). Anti-MAR detection (second panel) showed increase in MARTNKS (lanes 3, 4, and 5). A similar analysis using RNF114 and DTX2 in place of RNF166 and DTX3 yielded comparable results (Fig. 5C); RNF114 (lane 3) or DTX2 (lane 4) increased TNKS, and their combination led to a greater increase (lane 5).

We assessed the impact of the DTX3/RNF166 pathway on endogenous TNKS in the presence of RNF146. MycRNF146 was transfected into HEK293T WT cells, and the effects of Di19UIM and DTX3 were analyzed (Fig. 5D). Both Di19-UIM (lane 3) or DTX3 (lane 4) increased TNKS levels, with their combination resulting in the greatest increase (lane 5). Similarly, in the case of RNF166 and DTX3 (Fig. 5E), RNF166 (lane 3) or DTX3 (lane 4) increased TNKS levels, with their combination showing the highest increase (lane 5).

### DTX3 is monoubiquitylated on MAR

We investigated whether proteins other than tankyrase were impacted by the DTX2/3- RNF114/166 pathway described above. During our experiments, we detected (what could be) ubiquitylated DTX2 or DTX3 following immunoisolation of HA-ubiquitylated TNKS SAMPARP (see Fig. 2B, G, J, K for DTX2, and Fig. 3D, H for DTX3). This ubiquitylation likely resulted from auto-ubiquitylation as it was absent in catalytically inactive mutants (see Fig. 2G, lane 8 for DTX2 and Fig. 3D, lane 8 for DTX3). We focused on DTX3, which showed stronger activity than DTX2 in our assays.

To determine if DTX3 ubiquitylation was independent of TNKS SAMPARP, we introduced HA-Ub and FlagDTX3 or FlagDTX.CD (the catalytically dead mutant) into TNKS1/2 DKO cells, immunoisolated HA-ubiquitylated proteins, and analyzed by Flag immunoblot. Monoubiquitylated DTX3 was detected (Fig. 6A, lane 5). We also observed unmodified DTX3, which could be due to dimerization of HA-Ub-DTX3 with DTX3 (dimerization has been described for DTX2^33^) or to degradation of the modification during isolation. DTX3.CD was not ubiquitylated (lane 6), consistent with an auto-ubiquitylation mechanism. Immunoblot with anti-MAR antibody showed that monoUbDTX3 (and DTX3) was MARylated (second panel, lane 5).

**Figure 6.**
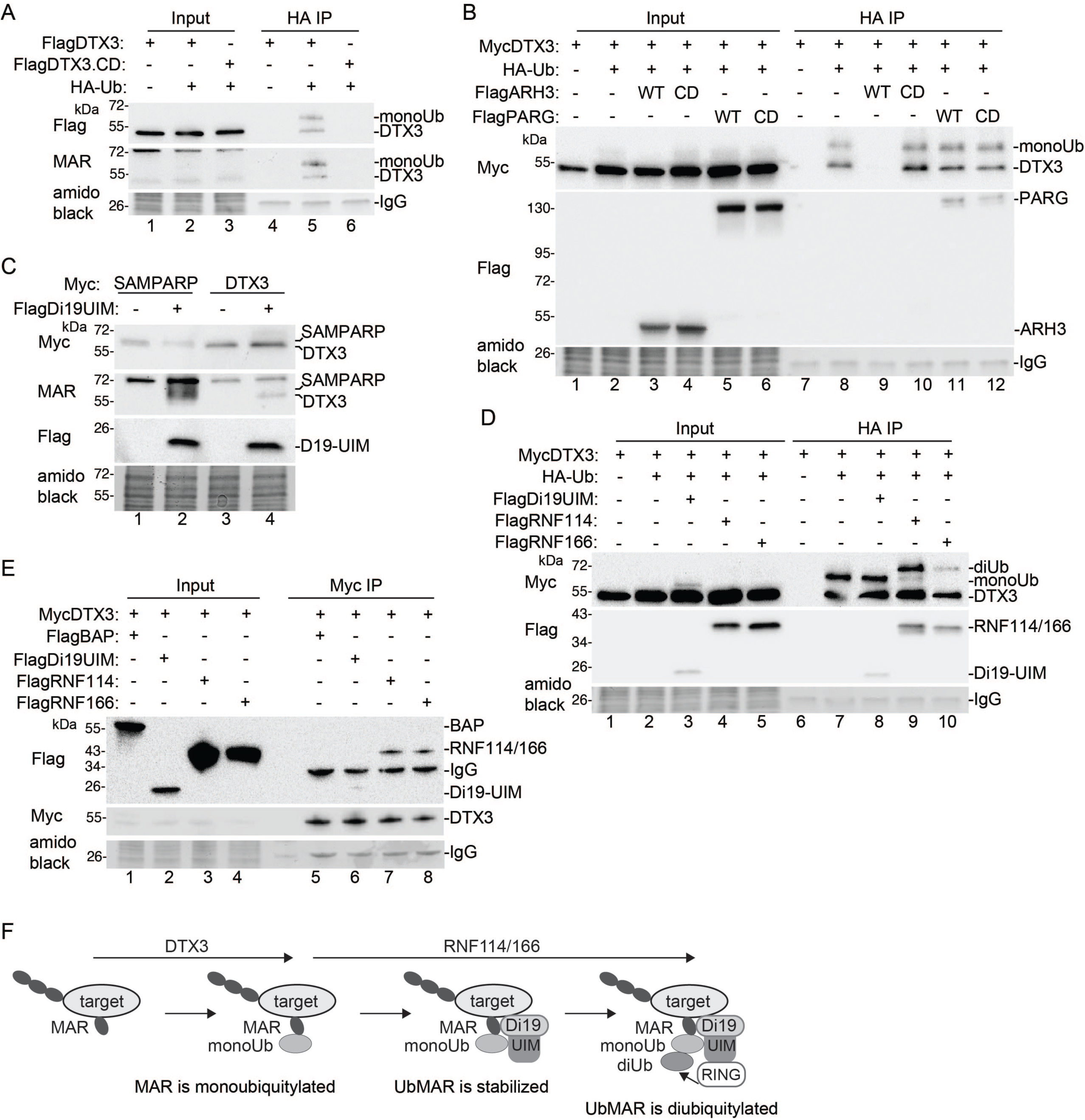
DTX3 is monoubiquitylated on MAR. B. Immunoblot analysis of TNKS1/2 DKO HEK293T cells transfected with the FLAGDTX3 or FLAGDTX3.CD and HA-Ub, immunoprecipitated with anti-HA antibody, stained with amido black, and probed with antibodies indicated on the left. C. Immunoblot analysis of TNKS1/2 DKO HEK293T cells transfected with the MycDTX3, HA-Ub and the indicated Flag plasmids, immunoprecipitated with anti-HA antibody, stained with amido black, and probed with antibodies indicated on the left. D. Immunoblot analysis of TNKS1/2 DKO HEK293T cells transfected with MycSAMPARP or MycDTX3, and FlagDi19UIM, stained with amido black, and probed with antibodies indicated on the left. E. Immunoblot analysis of TNKS1/2 DKO HEK293T cells transfected with the MycDTX3, HA-Ub and the indicated Flag plasmids, immunoprecipitated with anti-HA antibody, stained with amido black, and probed with antibodies indicated on the left. F. Immunoblot analysis of TNKS1/2 DKO HEK293T cells transfected with MycDTX3 and the indicated Flag plasmids, immunoprecipitated with anti-Myc antibody, stained with amido black, and probed with antibodies indicated on the left. 1. Schematic diagram showing a general mechanism where a MARylated protein target is monoubiquitylated on MAR by DTX3, followed by RNF114/166-mediated stabilization and dibuiquitylation.

We next asked if the monoubiquitin was on MAR on DTX3. We introduced HA-Ub, MycDTX3, and ARH3 (WT or CD) or PARG (WT or CD) into TNKS1/2 DKO cells, immunoisolated HA-ubiquitylated proteins, and analyzed by Myc immunoblot. Monoubiquitylated DTX3 was detected (Fig. 6B, lane 8). The DTX3 signal was eliminated by WT ARH3 (Fig. 6B, lane 9), but not by CD (lane 10), and was unaffected by PARG WT (lane 11) or CD (lane 12), indicating that DTX3 auto-monoubiquitylation occurs on a MARylated amino acid (likely serine).

Since DTX3 is monoubiquitylated on MAR, like TNKS SAMPARP, we asked if Di19-UIM affects the MAR modification on DTX3, as it does on TNKS SAMPARP. TNKS1/2 DKO cells were transfected with MycDTX3 (or MycSAMPARP as control) and FlagDi19UIM and analyzed by immunoblot. As shown in Fig. 6C, Di19-UIM induced MARDTX3 (lane 4). Next, we asked if the RNF114/166 RING domain catalyzes diubiquitylation of DTX3, as it does on TNKS SAMPARP. We introduced DTX3 and HA-Ub into TNKS1/2 DKO cells, immunoisolated HA-ubiquitylated proteins, and analyzed by immunoblot. We detect monoubiquitylated DTX3, both with and without Di19UIM (Fig. 6D, lanes 7 and 8), as expected. Adding full-length RNF114 (lane 9) or RNF166 (lane 10), induced diubiquitylation of DTX3. Our observations, that Di19-UIM stabilizes MAR DTX3 and RNF114/166 diubiquitylate monoUbMARDTX3 in TNKS1/2 DKO cells, suggests that these E3 ligases (DTX3 and RNF114/166) interact in the absence of TNKS. Coimmunoprecipitation analysis confirmed that MycDTX3 interacts with Di19UIM, RNF114, and RNF166 in TNKS1/2 DKO cells (Fig. 6E, lanes 6, 7, and 8).

## Discussion

Tankyrases have a broad impact on protein function in cells. In addition to their ankyrin repeat domains, which bind over forty partners^3^, they have PTMs (MAR, PAR, and ubiquitin), which bind reader proteins. Here we reveal unexpected crosstalk between tankyrase, PTMs and E3 ligases that affect tankyrase stability and function. Tankyrase catalyzes formation of PAR chains, which are bound by the E3 ligase RNF146 to promote degradation. But PAR can also be modified by PARG to produce MAR. Tankyrase is modified by both MAR and PAR in cells^17^. Here we show that the distribution of MAR and PAR on tankyrase can be influenced by other E3 ligases. We showed that Deltex E3 ligases bind PARylated tankyrase and monoubiquitylate MAR, creating a hybrid mark. The RING-UIM E3 ligases RNF114 and RNF166 then bind monoubiquitylated MAR and add a second K11-linked ubiquitin. We propose that the initial ubiquitylation of MAR and the subsequent binding and addition of a second ubiquitin blocks PAR formation, antagonizing the action of the PAR-binding E3 ligase RNF146 and stabilizing tankyrase.

We identified the Deltex family of E3 ligases as responsible for monoubiquitylation of MAR on TNKS. The family includes five members: DTX1, 2, 3, 3L, and 4. All except DTX3L, bind TNKS dependent on TNKS catalytic activity. All members share a RING (catalytic) domain and CTD (ADP-ribose-binding) domain, but only DTX1, 2, and 4 have PAR-binding WWE domains in their N termini. Surprisingly, deleting the WWE domains (shown for DTX2) did not prevent binding or ubiquitylation of TNKS. The RING-DTC fragment (shown for DTX3) was sufficient for binding to PARylated TNKS and monoubiquitylating it on MAR. Whether the WWE domains impact DTX2 targets and function in cells remain to be determined.

The amino acid in TNKS that MAR is attached to is likely serine. While many studies have identified PARP1 ADP-ribosylation targets, less is known about tankyrase^34, 35^. Our data are most consistent with ubiquitylation of monoADP-ribose on serine. We tested this using the hydrolases ARH3 and PARG in cells and in vitro. ARH3 cleaves the O-glycosidic linkage of serine-ADP-ribose and, to a lesser extent, poly-ADP-ribose, whereas PARG primarily degrades poly-ADP-ribose but has limited activity on the ADP-ribose-protein bond^6^. In vitro assays showed that ARH3 removed monoubiquitin and MAR from TNKS SAMPARP, while PARG had no effect. This was confirmed in cells, where ARH3’s catalytic was required and PARG showed no effect.

Deltex E3 ligases specifically target the 3’hydroxyl of the ADP-ribosyl moiety that can be linked to a protein^24^. This site is near the 2’hydroxyl, where the next ADP-ribose unit is added during PAR chain synthesis. A prediction is that ubiquitylation of the 3’hydroxyl could block ADP-ribose addition at the adjacent 2’hydroxyl, limiting PAR and promoting MAR. Additionally, a protein with a single reader domain capable of binding to either ubiquitin or MAR could further inhibit PARylation. A hybrid reader, like Di19-UIM, which binds both ubiquitin and MAR, could enhance specificity and binding strength. Mutational analysis showed that disrupting either the Di19 (MAR-binding) or UIM (ubiquitin-binding) domains eliminated Di19-UIM binding and MAR stabilization. Binding of the Di19-UIM hybrid reader in the context of full length RNF114/166 positions the N-terminal RING domain to cap the hybrid ubiquitin-MAR mark with a second (K11- linked) ubiquitin. This process, where the Deltex E3 ligase creates the hybrid mark, coupled with hybrid reader binding and further modification by a second E3 ligase RNF114/166, dramatically impacts ADP-ribosylation and subsequent binding to a third E3 ligase (RNF146).

Tankyrase is not the only target of this tandem E3 ligase pathway. We showed that DTX3 is also monoubiquitylated on MAR, likely on serine, as the Ub-MAR hybrid mark was cleaved by ARH3. Like TNKS SAMPARP, MARylation on monoUbDTX3 was stabilized by the RNF166 Di19-UIM domain, with a second ubiquitin added by the RNF114/166 amino terminal domain. Thus, DTX3 and RNF114/166 collaborate; where DTX3 adds the hybrid mark (to itself or TNKS) and RNF114/166 reads and further modifies it (Fig. 6F). Interestingly, a recent study identified PARP10 as being ubiquitylated on MAR and extended with K11-linked polyubiquitin chains^36^. Whether this relies on the same E3 ligases described here remains to be determined, but it nonetheless highlights a growing number of examples of ADP-ribosylation/ubiquitylation crosstalk in cells. Considering their inherent power and specificity, we anticipate that future studies will reveal new hybrid marks and hybrid readers that underlie complex signaling pathways in cells.

## Material and Methods

### Cell lines

HEK293T (ATCC) and HEK293T TNKS1/2 DKO^25^ cell lines were supplemented with 10% DBS and grown in standard conditions.

### Cell transfection and treatment

Cells were seeded in 6 well plates and treated the next day using 2 µg of plasmid and 4 µL of lipofectamine 3000 (Invitrogen) for 24 h according to the manufacturer’s instructions. Cells were harvested with trypsin and washed in cold PBS. Tankyrase inhibitor #8 (TNKSi)^37^ was used at 10 µM for 24 h (Chembridge Corporation, MolPort-000-222-699).

For siRNAs, cells were transfected with Lipofectamine RNAiMAX (Invitrogen) according to the manufacturer’s protocol for 48-72 hr. The final concentration of siRNA was 20 nM. The following target sequence was used for DTX2 siRNA: DTX2-2 (5’- CCUCAUAGUUUACAGCAUU- 3’)^38^ (Dharmacon). The control siRNA is the GFP duplex I (Dharmacon)

### Plasmids

The following plasmids were provided: pCMV-3XF-RNF166^39^(by Ramnik Xavier); pcDNA3-Myc- DTX1, DTX2, DTX3, DTX3L, and DTX4^40^ (by Danny Huang); and EGFP-N1-hPARGflwt (by Francois Dantzer) or obtained from Addgene: pcDNA3.1-RNF114-myc-His (58295, from Francesca Capon); pGEX6P-2-hRNF146-FL (132610, from Wenqing Xu); pRK5-HA-Ubiquitin WT (17608, from Ted Dawson); and pET30-ARH3-His-Sumo-HA (111578, from Thomas Muir). pLPCFlagTNKS1^41^; TNKS.WT (TT20.WT)^42^; and 3XFlag-BAP (Sigma) were described previously. The following plasmids were described previously: pCMV3FlagDi19UIM; pCMV3FlagSAMPARP; pCMV3FlagRNF114; pCMV3MycDi19UIM; pCMV3MycSAMPARP, pCMV3MycRNF166; pRK5-3FlagDTX2; and pRK5-3MycDTX2^17^. The following plasmids were generated: pCMV3FlagSAMPARP.EA (E1291A, based on the PARP1 mutant E998A^32^); pCMV3MycRNF114 (1 to 228); pCMV3MycDi19UIM.Mut1(C181G^17^); pCMV3MycDi19UIM.Mut2 (A229P/S233Q^17^); pRK5-3MycDTX3.WT (1 to 347); and pRK5-3MycRNF146.WT (1 to 359).

The following DTX2 plasmids were generated in pRK5-3Flag: FlagDTX2.WT (1 to 622); FlagDTX2.CD (C412A/C415A, based on the DTX3L mutant C561S/C564S^31^); FlagDTX2.WWE (1 to 187); FlagDTX2.RD (384 to 622); FlagDTX2.WWE** (W24A/W114A^14, 23^); and FlagDTX2.PRD (176 to 622). The following DTX3 plasmids were generated in pRK5-3Flag: FlagDTX3.WT (1 to 347); FlagDTX3.CD (C164S/C167S, based on the DTX3L mutant C561S/C564S^31^); FlagDTX3.Pro (1 to 165); and FlagDTX3.RD (152 to 347). The DTX3L plasmids FlagDTX3L.WT (1 to 740) and FlagDTX3L.RD (545 to 740), the ARH3 plasmids FlagARH3.WT (13 to 463) and FlagARH3.CD (D77N/D78N^28^), and the PARG plasmids FlagPARG.WT (1 to 976) and FlagPARG.CD (E755A/E756A^29^) were generated in pRK5-3Flag.

Plasmid DNA sequences were modified using PCR amplification, enzymatic restriction, and ligation, or site-directed mutagenesis (Agilent) and (NEB) or DNA assembly (NEBuilder® HiFi DNA Assembly Master Mix).

### Protein extraction and immunoprecipitation

Proteins were extracted by resuspending the cell pellets for 1 h on ice in TNE buffer [10 mM Tris (pH 7.8), 1% Nonidet P-40, 0.15 M NaCl, 1 mM EDTA, 2.5% protease inhibitor cocktail (Sigma, P8340), 1 µM of PARGi (Sigma), and 20 mM NEM (Sigma)]. NEM was omitted for the experiment in Fig. 1B. The lysates were pelleted at 10,000 g for 10 min and the supernatants were used to determine the protein concentration using Bradford assay (Bio-Rad). After a preclearing step with Protein G Sepharose (Sigma), equal amounts of proteins were then incubated with anti-Flag Beads (Sigma), anti-Myc Beads (Sigma) or anti-HA Beads (Sigma) for 2 h at 4°C under agitation. The beads were washed at least three times with 1 mL of TNE buffer. The samples were denatured in Laemmli buffer at 70°C for 10 min.

Immunoprecipitations in Figures 1E, 2B, and 3G, were performed under denaturing conditions. Pelleted cells were lysed using SDS lysis buffer (2% SDS, 0.9% Nonidet P-40, 9mM Tris pH 8.0, 135 mM NaCl, 0.9 mM EDTA) and heated at 70°C for 10 min. After a ten-fold dilution with TNE buffer, samples were sonicated for 1 min using a Diagenode bioruptor device and then centrifuged 21,000 g for 10 min at 4°C. Supernatants were incubated with anti-HA beads for 2 h. The beads were then washed three times with TNE and one time with 1M NaCl lysis buffer (20 mM Tris pH 8.0, 1M NaCl, 1% Triton X-100, 2 mM EDTA, 0.2% SDS). Samples were resuspended in Laemmli buffer and heated at 70°C for 10 min.

### Treatments in vitro

Reactions were performed following the last wash of the HA-IP experiment. For ARH3 or PARG treatment, 10 µL of beads were incubated in 30 µL PARP buffer (4 mM MgCl2, 50 mM Tris pH 8.0, 0.2 mM DTT) with 1 µM ARH3 (NKmax, ATGP1665) or 200 ng of PARG (AdipoGen, AG-40T- 0022). For Cezanne treatment, 10 µL of beads were incubated in 30 µL of DUB reaction buffer (50 mM Tris pH 7.5, 50 mM NaCl, 5 mM DTT) with 0.2 µM Cezanne (UbpBio, H4200)^26^. For NH2OH treatment, 10 µL of beads were incubated in 30µL reaction buffer [50 mM HEPES (pH 7.5), 50 mM NaCl, 5 mM MgCl2, and 1 mM DTT] with 400 mM NH2OH (Fisher, AC270101000)^24^. Water was used for control treatments. After 30 minutes at 37°C under shaking at 1000 rpm, the reactions were stopped by addition of 10 µL 4X Laemmli buffer and denatured at 70°C for 10 min. For NH2OH treated samples, the pH was adjusted by adding 1 µL of 30% NaOH.

### Immunoblot analysis

Protein samples were loaded using precast gels (Bio-Rad), subjected to SDS-PAGE, and transferred onto nitrocellulose membrane using wet transfer of 100V for 1h. The membranes were stained with amido black and incubated with the following primary antibodies: rabbit anti-TNKS 762^43^; rabbit anti-TNKS763^43^; mouse anti-Flag M2 F3165 (Sigma); mouse anti-gamma-tubulin T5168 (Sigma); mouse anti-Myc 4A6 05-724 (Millipore); rabbit anti-RNF146 PA5-55544 (Invitrogen); rabbit anti-HA ab9110 (Abcam); rabbit anti-RNF114 HPA021184 (Sigma); or rabbit anti-DTX2 PA5-60164 (ThermoFisher), followed by HRP coupled secondary antibodies. For detecting PAR, the anti-poly-ADP-ribose binding reagent MABE1031 (Sigma) was used. For detecting mono-ADP-ribose, anti-MAR antibody (AbD43647) BioRad was incubated with BiSpyCatcher2-HRP (TZC002P) BioRad at a ratio 10:0.8 (v/v) for 1 hour before a 5000-fold dilution in 5% non-fat milk PBST. The signal was acquired using ECL (Fisher) and the ChemiDoc MP imaging system (Bio-Rad).

### Statistical analysis

Statistical analysis was performed using Prism 10 software. Data are shown as mean ± SEM. Student unpaired *t* test was applied. P < 0.05 values were considered significant: *, P ≤ 0.05; **, P ≤ 0.01.

## Acknowledgements

We thank Francois Dantzer, Danny Huang, and Ramnik Xavier for plasmids. Research reported in this publication was supported by the National Institute of General Medicine Sciences of the National Institutes of Health under award numbers R35GM149355 and R01GM141292 to S.S.

## Author contribution

J.P., K.G., K.R., and S.S. conceived the experimental design, analyzed the data, and wrote the manuscript. J.P., K.G., K.R. performed all experiments.

## Notes

### Competing Interest Statement

The authors have declared no competing interest.

